# Metabolomic changes associated with acquired resistance to *Ixodes scapularis*

**DOI:** 10.1101/2023.07.31.551287

**Authors:** Yingjun Cui, Jaqueline Matias, Xiaotian Tang, Balasubramanian Cibichakravarthy, Kathleen DePonte, Ming-Jie Wu, Erol Fikrig

**Author notes:** Corresponding authors: Yingjun Cui and Erol Fikrig. Section of Infectious Diseases, Department of Internal Medicine. Yale University School of Medicine. Room 169, 300 Cedar Street. New Haven, Connecticut 06520-8031. and.

## Abstract

Guinea pigs repeatedly exposed to *Ixodes scapularis* develop acquired resistance to the ticks (ATR). The molecular mechanisms of ATR have not been fully elucidated, and partially involve immune responses to proteins in tick saliva. In this study, we examined the metabolome of sera of guinea pigs during the development of ATR. Induction of components of the tyrosine metabolic pathway, including hydroxyphenyllactic acid (HPLA), were associated with ATR. We therefore administered HPLA to mice, an animal that does not develop ATR, and exposed the animals to *I. scapularis*. We also administered nitisinone, a known inhibitor of tyrosine degradation, to another group of mice. The mortality of *I. scapularis* that fed on mice given HPLA or nitisinone was 26% and 72% respectively, compared with 2% mortality among ticks that fed on control animals. These data indicate that metabolic changes that occur after tick bites contribute to ATR.

## Introduction

Ticks are an important medical and veterinary global public health concern due to direct damage caused by blood feeding, and their roles in transmitting numerous infectious agents including bacteria, viruses, and protozoa(*1-3*). Pathogens transmitted by ticks are responsible for the majority of vector-borne diseases in temperate North America and Europe (*3*). In North America, *Ixodes scapularis* is common and transmits *Borrelia, Anaplasma, Babesia* and *Powassan* virus, among other pathogens. The geographic range of ticks is rapidly expanding with climate change and the number of cases of *I. scapularis*-borne diseases is similarly increasing (*3-6*). Therefore, new strategies to prevent or decrease *I. scapularis*-borne illnesses are needed (*4, 7*).

Guinea pigs, cattle and other animals can develop acquired tick resistance (ATR), which is manifested by increased redness at the tick bite site, altered tick attachment, decreased tick engorgement, reduced molting capacity and fecundity, increased tick mortality, and in some cases, an alteration in the transmission of specific tick-borne pathogens (*8-17*). Tick feeding induces host immune regulatory and effector pathways, resulting in tick-specific antibody production, complement activation, and cellular immune responses (*9, 10, 12*). In addition, bioreactive molecules released by basophils, eosinophils, and mast cells may influence tick physiology (*9*). Histamine is an effector of immediate cutaneous hypersensitivity reactions, and can have a direct effect on tick attachment, and the development of ATR (*18, 19*).

Metabolites drive essential cellular functions, including energy production and storage, signal transduction and apoptosis (*20*). Metabolomics is defined as the comprehensive analysis of metabolites in a biological specimen. An analysis of metabolites directly reflects the underlying biochemical activity and state of cells and tissues, and can help to define a molecular phenotype (*21*). In this study, we investigated the metabolome of guinea pigs before and after tick bites, and during the development of ATR, in order to characterize metabolic pathways that are associated with ATR.

## Results

Upon repeated exposure to *I. scapularis*, guinea pigs develop ATR. Studies were therefore performed to examine metabolic changes in guinea pigs that may be associated with ATR. First, guinea pigs were exposed to *I. scapularis* to generate ATR. In previous studies, the placement of 30 ticks, twice at two-week intervals has been commonly used to induce ATR. Here we now show that the placement of 3 ticks on guinea pigs is sufficient to elicit ATR. Three nymphal *I. scapularis* were used for the first challenge in five guinea pigs, and 25 nymph ticks were used for second challenge to better examine how rapidly a large group of ticks were affected. After the second tick challenge, all five guinea pigs developed erythema at tick bite sites (Fig. 1A). Moreover, the mean tick attachment rates (73.6%, 61.6% and 38.4%) at 24, 48 and 72 hours were significantly decreased compared to first tick challenge (100%, 100% and 100%) (Fig.1B). The mean tick weight (1.1 mg) was also significantly lower than following first tick challenge (2.8 mg) (Fig.1C).

**Figure. 1.**
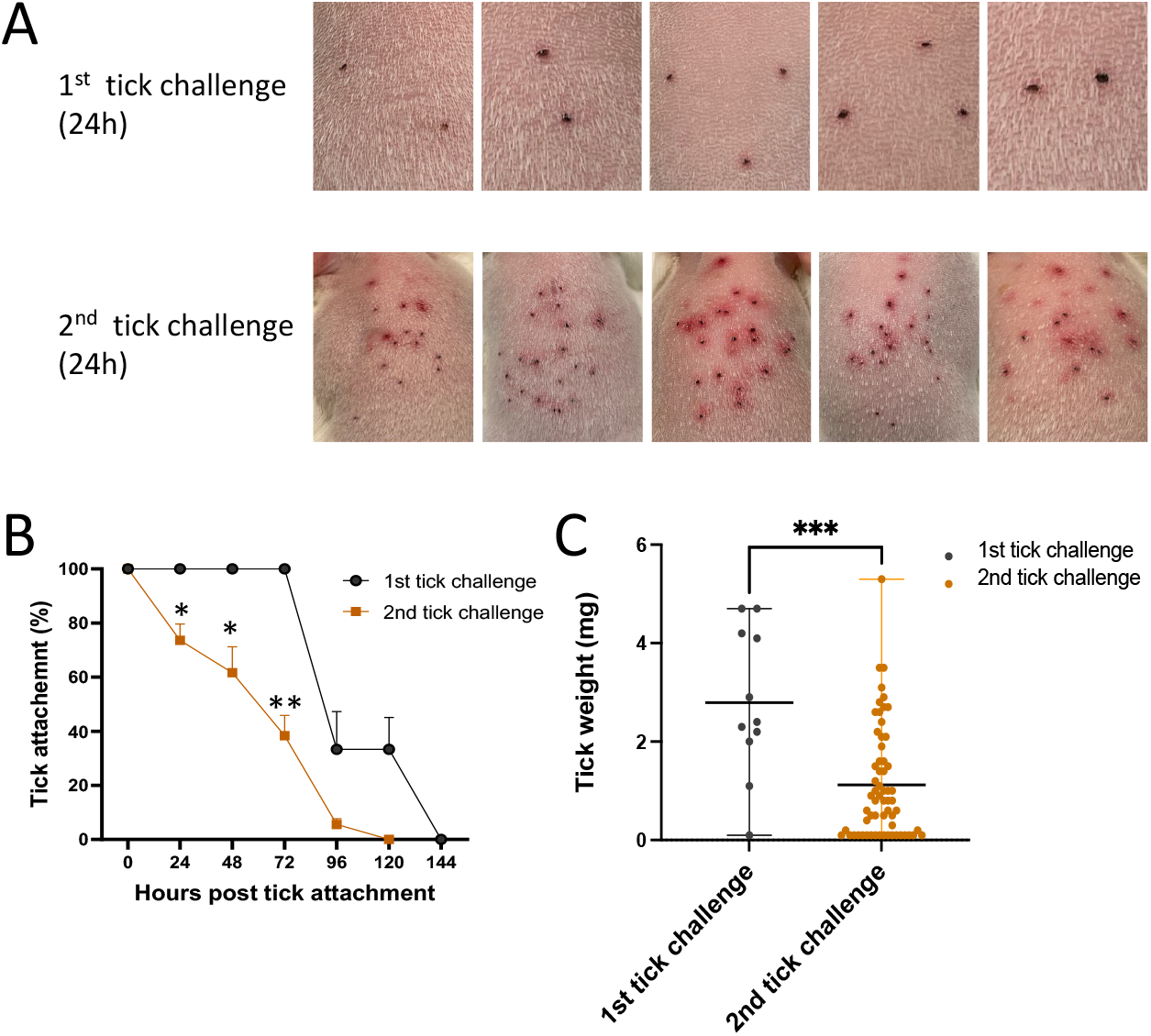
Guinea pigs develop acquired tick resistance after exposure to 3 ticks. **(A)** Images of a guinea pig at first and second tick challenge (24 h post tick challenge). **(B)** Tick attachment rates (%) in guinea pigs at first and second tick challenge. **(C)** Weights of the fed nymphal ticks at first and second tick challenge. The individual weights and means with range are represented. The * means p value<0.05, ** means p value<0.01, ***means p value<0.001.

In contrast to guinea pigs, mice do not develop ATR following exposure to *I. scapularis*. Mice can therefore serve as an additional comparison with guinea pigs. As expected, mice did not develop ATR when exposed to 10 *I. scapularis*, 3 times at 2-week intervals. There was no alteration in erythema at the tick bite sites (Fig. 2A), tick attachment rates (Fig. 2B), or tick weights (Fig. 2C).

**Figure. 2.**
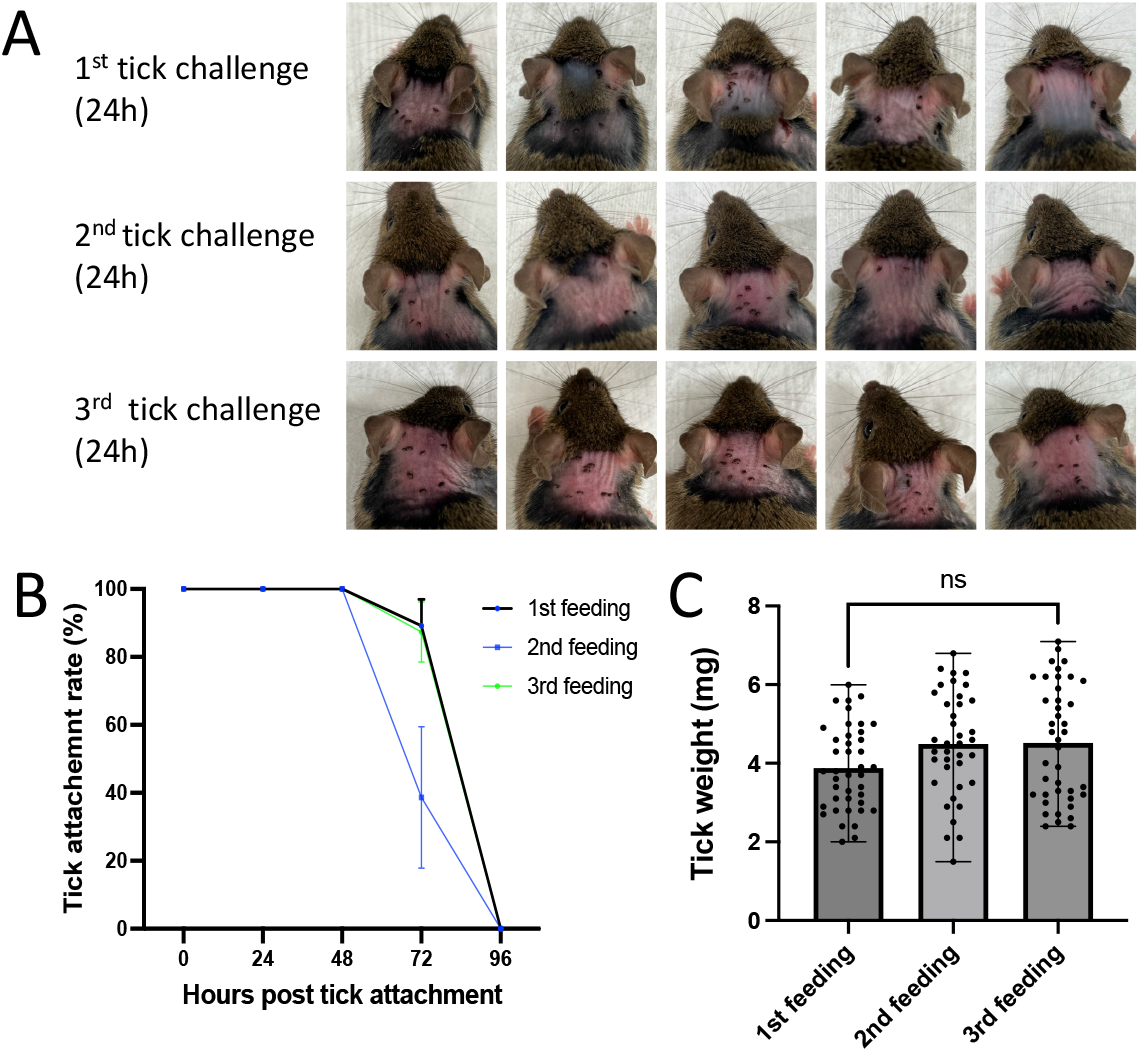
Mice do not develop acquired resistance to ticks. **(A)** Images of mice at first, second and third tick challenge (24 h post tick challenge). **(B)** Tick attachment rate (%) in mice at the first, second tick and third tick challenge. **(C)** Weights of the fed ticks. The individual weights and means with range are represented.

UPLC–MS was performed with sera from guinea pigs before and after tick challenge. Both ESI- and ESI+ were used with the data from UPLC–MS. PCA was used to compare metabolomes of guinea pigs before and after tick challenge. The results indicated that the metabolomes from guinea pig after tick challenge clustered closely and separated from the metabolomes of guinea pig before tick challenge (Fig. 3A, B). 187 metabolites were detected in the sera of guinea pigs in ESI-mode, 49 metabolites were significantly up-regulated (variable importance in the projection (VIP) value >1.5, fold change>2, and p value<0.05), and 17 metabolites were significantly down-regulated (VIP value >1.5, fold change<0.5, and p value<0.05) (Fig. 3C, supplemental table 1).

**Figure. 3.**
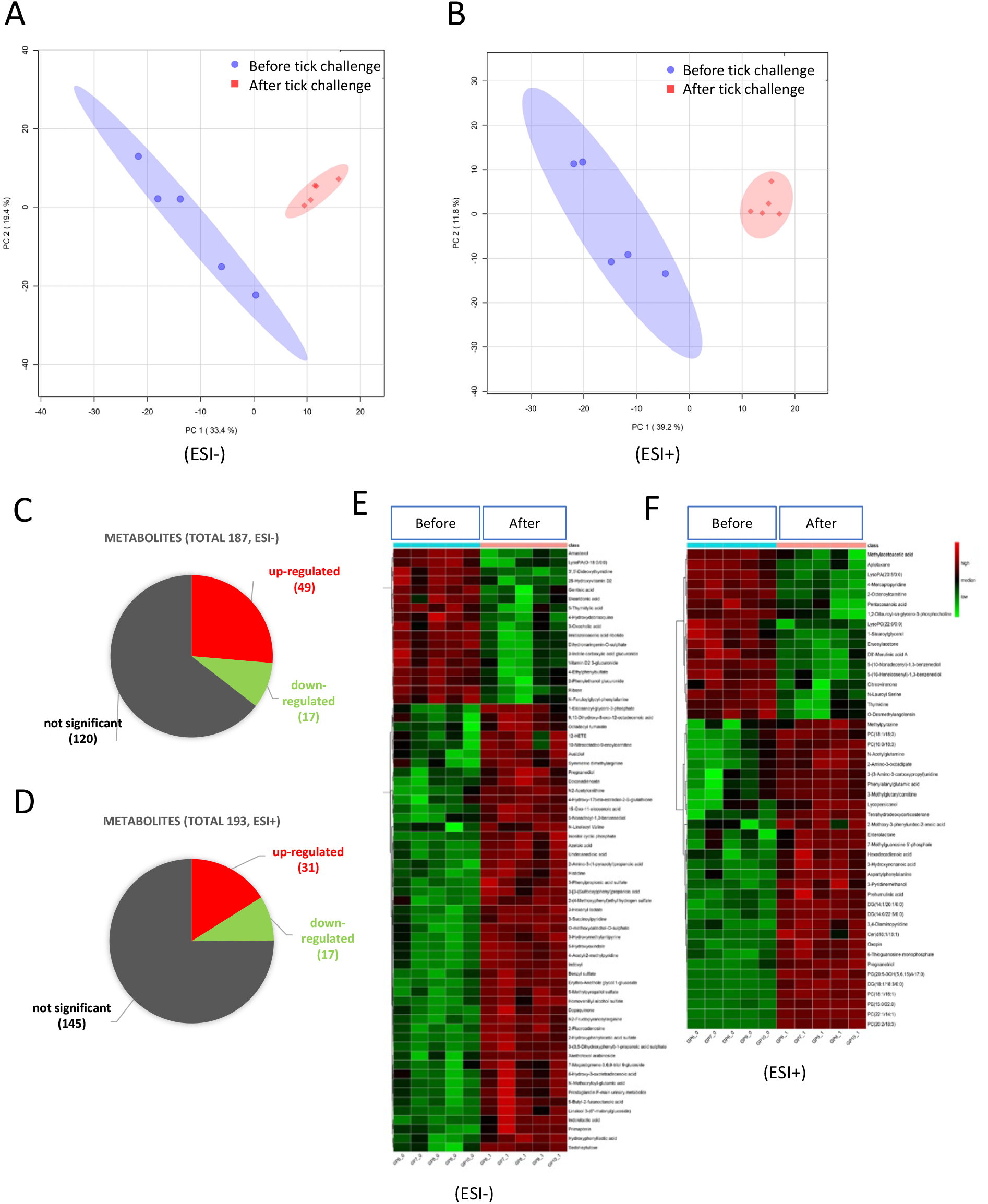
Metabolic change in guinea pigs that are associated with acquired tick resistance. The UPLC–MS was performed in the untargeted metabolomics analysis. Both ESI+ and ESI-were used for mass scan in UPLC–MS. **(A)** and **(B)** Principal component analysis of the metabolomes of guinea pigs before and after tick challenge. C and D. Summary of the detected metabolites and differentially expressed metabolites in guinea pigs. E and F. Heatmap showing expression of the differentially expressed metabolites.

193 metabolites were detected from sera of guinea pigs in ESI+ mode, 31 metabolites were significantly up-regulated (VIP value >1.5, fold change>2, and p value<0.05), and 17 metabolites were significantly down-regulated (VIP value >1.5, fold change<0.5, and p value<0.05) (Fig.3D, supplemental table 2). The heatmap showing expression of differentially expressed metabolites also indicated that there is a distinct expression pattern between samples from guinea pigs before and after tick challenge, and there is highly similarity between samples within same group. As mice do not develop ATR, we also characterize the metabolomes of mice upon three *I. scapularis* challenges and compared this to the metabolome of mice without tick challenge. PCA results indicated that the metabolomes from tick-challenged mice were clearly separated from control mice (Fig. 4A, B). 204 metabolites were detected from sera of mice in ESI-mode, 23 metabolites were significantly up-regulated (VIP value >1.5, fold change>2, and p value<0.05), and 29 metabolites were significantly down-regulated (VIP value >1.5, fold change<0.5, and p value<0.05) (Fig.4C, supplemental table 3). 194 metabolites were detected from sera of mice in ESI+ mode, 14 metabolites were significantly up-regulated (VIP value >1.5, fold change>2, and p value<0.05), and 44 metabolites were significantly down-regulated (VIP value >1.5, fold change<0.5, and p value<0.05) (Fig.4D, supplemental table 4). The heatmap showing expression of differentially expressed metabolites also indicated that there is a clear distinct expression pattern between samples from mice with and without tick challenge (Fig.3E, F). These data also indicate that tick bites changed the metabolomes of mice upon three tick challenges.

**Figure. 4.**
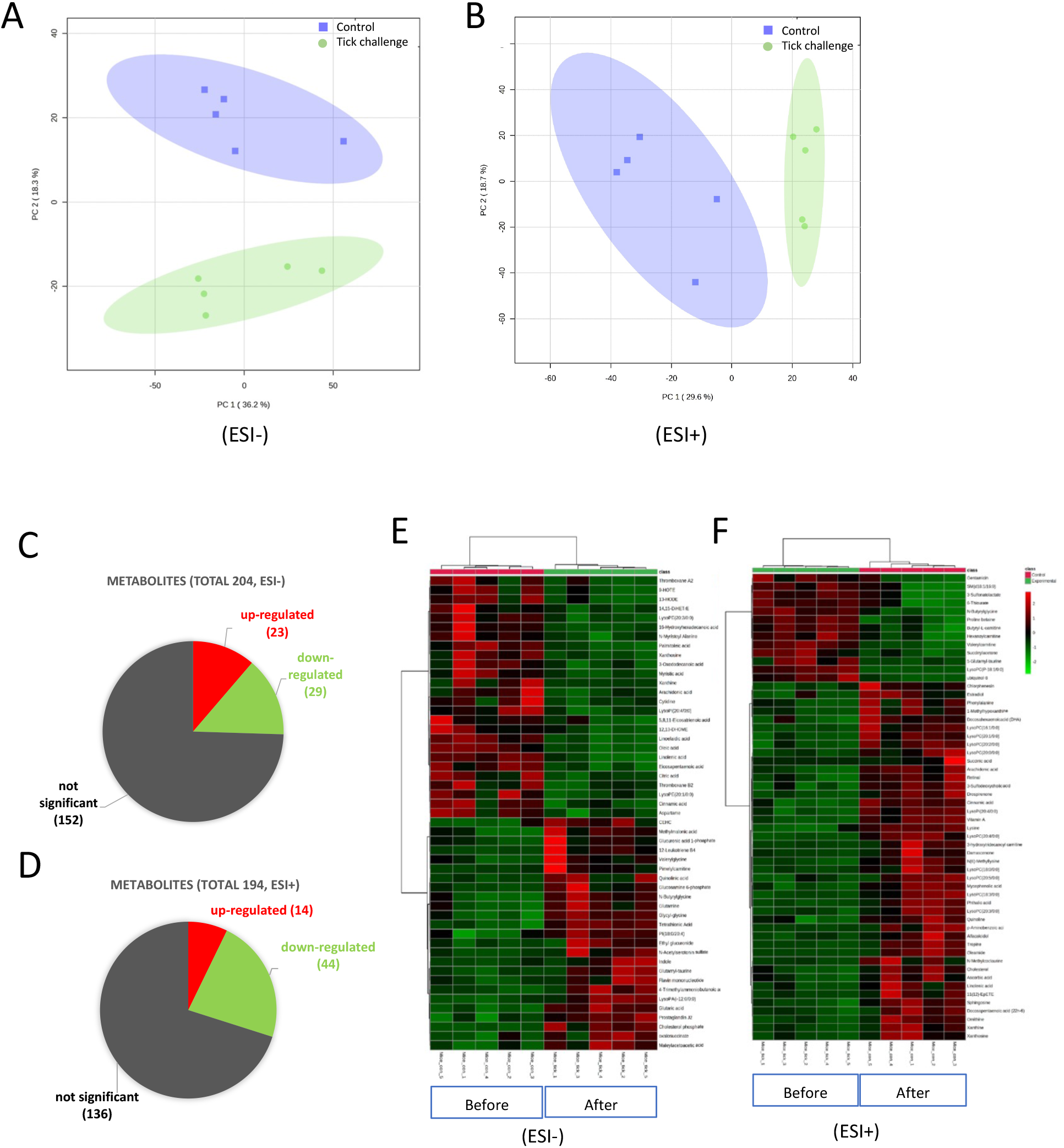
Metabolomic changes in mice upon tick exposure. UPLC–MS was performed in the untargeted metabolomics analysis. Both ESI+ and ESI-were used for mass scan in UPLC–MS. **(A)** and **(B)** Principal component analysis of the metabolomes of mice with and without tick challenge. **(C)** and **(D)** Summary of the detected metabolites and differentially expressed metabolites. E and F. Heatmap showing expression of the differentially expressed metabolites.

The differentially expressed metabolites from the ESI-mode and ESI+ mode were combined and used for KEGG pathway analysis. Two pathways including tyrosine and pyrimidine metabolism were significantly enriched in guinea pigs (p value<0.05) (Fig.5A). Dopaquinone and hydroxyphenyllactic acid (HPLA) in tyrosine metabolism were significantly upregulated by 2.4 and 2.8-fold after tick bite, respectively. Thymidine, 5-thymidylic acid and deoxyuridine 5’-phosphate (dUMP) in pyrimidine metabolism were downregulated by 3.7, 4.8 and 6-fold after tick bite, respectively. In mice, alpha linolenic acid and linoleic acid metabolism, and carnitine synthesis are significantly enriched after tick bite (p value<0.05). Eicosapentaenoic acid, arachidonic acid, docosapentaenoic acid and linolenic acid, which are involved in alpha linolenic acid and linoleic acid metabolism were downregulated by 2.1, 2.4, 2.6 and 2.7-fold after tick bites.

**Figure. 5.**
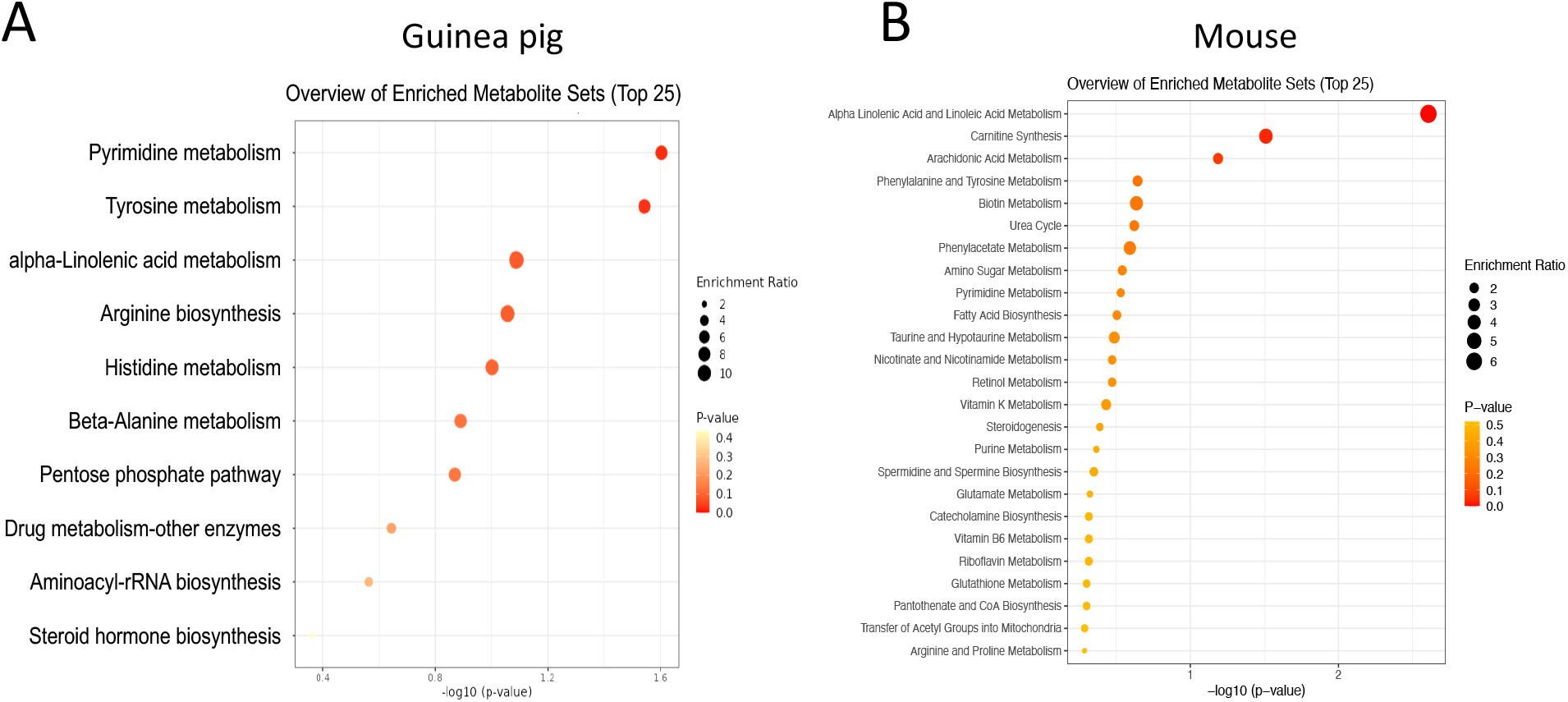
Tick exposure influences different metabolic pathways in guinea pigs and mice. **(A)** The enriched KEGG pathways for differentially expressed metabolites in guinea pigs. **(B)** The enriched KEGG pathways for differentially expressed metabolites in mice.

The inhibition of tyrosine degradation can injure bloodsucking arthropods including kissing bug, yellow fever mosquito and the Asian blue tick (*22*). Since components of the tyrosine metabolism pathway, including HPLA, were significantly enriched in guinea pigs with ATR, we tested whether administrating HPLA or providing a known inhibitor of tyrosine degradation, nitisinone, affected tick survival or tick rejection. We administered these compounds to mice, which do not develop ATR. Mice were given HPLA (1 mg/kg) or nitisinone (1 mg/kg) and *I. scapularis* were allowed to engorge on these animals. The ticks that fed on the HPLA-administered group (26.2%) or the nitisinone-administered group (71.8%) had higher death rates compared to ticks that fed on control mice (2.5%) (Fig.6). These data suggest that tyrosine metabolism affects *I. scapularis* survival following engorgement but does not influence the ability of ticks to feed on mice.

**Figure. 6.**
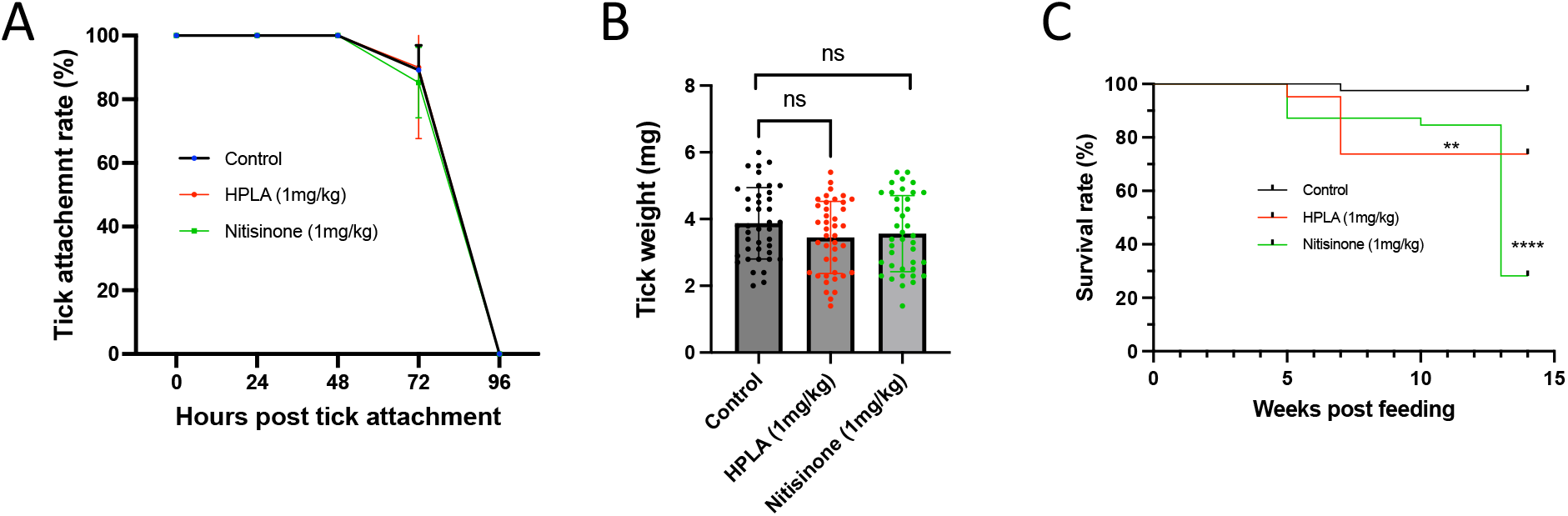
Targeting the tyrosine metabolic pathway in mice causes increased mortality in ticks following engorgement. Mice were administered HPLA (1 mg/kg body weight), nitisinone (1 mg/kg body weight) or PBS (as control). Nymphal ticks were then allowed to feed on these mice. **(A)** Feeding HPLA or nitisinone to mice did not affects tick attachment. (B) Feeding HPLA or nitisinone to mice did not affects weight of fed ticks. **(C)** Feeding HPLA or nitisinone to mice significantly decreased the survival rates of the fed tick. The log-rank (Mantel-Cox) test was used to assess survival curves, ** means p value<0.01, ****means p value<0.0001, ns means not significant.

## Discussion

*I. scapularis*-transmitted infectious agents are an increasing public health concern. Targeting tick feeding and interrupting its lifecycle, is one strategy for disease prevention or reduction (*23*). An anti-tick vaccine that mimics natural ATR is a promising approach, but progress is slow, in part because the mechanisms that contribute to ATR are not fully understood (*23*). An mRNA vaccine consisting of a lipid nanoparticle containing nucleoside-modified mRNAs encoding 19 *I. scapularis* salivary proteins, recapitulates many aspects of ATR in guinea pigs and can prevent transmission of the Lyme disease agent (*24*). Further work has delineated the relative contribution of each of the 19 components within 19ISP in ATR, but the mechanisms associated with ATR need to be elucidated (*25*).

Antibodies directed at tick proteins are thought to partially contribute to ATR but are not sufficient to independently recapitulate ATR (*1, 9*). Histamine produced by basophils at the tick bite site is associated with ATR (*19, 26, 27*). In addition, serotonin controls feeding in ants (*28*), blow flies (*29*), mosquitoes (*30*), and honeybees (*31*), and symbiont-regulated serotonin biosynthesis modulates tick feeding (*32*). Furthermore, inhibition of tyrosine degradation selectively kills hematophagous arthropods (*22*). These studies with small molecules prompted us to hypothesize that tick feeding stimulates host metabolic and physiologic pathways which participate in the development of some aspects of ATR.

The metabolomic studies demonstrate that guinea pigs with ATR have a significant enrichment in several pathways, and the tyrosine and pyrimidine pathways are prominent. Repeated tick bites also changed the metabolomes of mice, but the pathways were different that those that tick bites induced in guinea pigs. As mice do not develop ATR, we focused on the pathways that were selectively induced in guinea pigs with ATR. Furthermore, it has reported that blood-feeding arthropods produce large amounts of tyrosine and that inhibition of tyrosine degradation is detrimental to some arthropods (*22*). Since the tyrosine pathway was induced in guinea pigs with ATR, we determined whether enhancement of this pathway could influence tick survival in mice which do not develop ATR. Interestingly, ticks which fed on mice given HPLA or nitisinone, each of which should increase the concentrations of HPLA and/or tyrosine in murine serum, had significantly increased mortality compared with ticks that feed on control animals. These studies offer a “proof of principle” that metabolites contribute to successful tick feeding and survival.

The genesis of ATR is multifactorial. Diverse immune, physiologic and metabolic responses collectively contribute to ATR. These studies demonstrate that the metabolome of guinea pigs that have ATR is different than the metabolome of guinea pigs that have not been exposed to ticks, or that of mice that do not develop ATR. It is most likely that several metabolites contribute to successful tick feeding and survival, and that the tyrosine pathway is not unique. A better understanding of the how different metabolic pathways contribute to ATR will help lead to new ways to induce ATR, interfere with the tick life cycle, and potentially prevent pathogen transmission.

## Materials and Methods

### Ethics statement

All the animal work was done in accordance with the Guide for the Care and Use of Laboratory Animals (National Institutes of Health, USA). The procedures were approved by Yale University Institutional Animal Care and Use Committee.

### Animals and ticks

Female Hartley guinea pigs 6–8-week-old, and 6-week-old female C3H/HeN mice were purchased from Charles River Laboratories. The animals were maintained in a facility at Yale University. *I. scapularis* larvae were obtained from the Oklahoma State University. Larvae molted into nymph after feeding on C3H/HeN mice. Ticks were maintained at 85% relative humidity with a 14h light and 10h dark period at 23°C.

### Tick challenge

Guinea pigs were anesthetized with ketamine and xylazine, 3 nymphal ticks were placed on the shaved back of one group of guinea pigs (n=5) at first tick challenge. 25-30 nymphal ticks were placed on these guinea pigs for the second tick challenge. Erythema, tick attachment and tick weight were recorded every 24 hours. Blood samples were collected from each guinea pig via retro-orbital bleeding one week before first tick challenge and three weeks after the first tick challenge.

Mice were anesthetized with ketamine/xylazine, 10 nymphal ticks were placed on the shaved back of one group of mice (n=5). Three challenges were performed in the experimental group of mice. Another group of mice (n=5), not exposed to ticks, was used as control. Blood samples were collected from each mouse after the third tick challenge and collected from each control mouse at same time.

### Untargeted metabolomics

100 μL of sera from blood collected from guinea pigs and mice at different intervals before and after tick feeding were used for untargeted metabolomics analysis. The ultra-performance liquid chromatography mass spectrometry (UPLC–MS) was performed in the untargeted metabolomics analysis. Separation was performed by ACQUITY UPLC (Waters) combined with Q Exactive MS (Thermo) and screened with ESI-MS. The LC system is comprised of an ACQUITY UPLC HSS T3 (100×2.1mm×1.8 μm) with ACQUITY UPLC(Waters). The mobile phase is composed of solvent A (0.05% formic acid water) and solvent B (acetonitrile) with a gradient elution (0-1 min, 5% B; 1-12 min, 5%-95% B; 12-13.5 min, 95% B; 13.5-13.6 min, 95%-5% B; 13.6-16 min, 5% B). The flow rate of the mobile phase is 0.3 mL/min. The column temperature is maintained at 40°C, and the sample manager temperature is set at 4°C. Both positive ion mode (ESI+) and negative ion mode (ESI-) were used for mass scan. The raw data were acquired and aligned using the Compound Discover (3.0, Thermo) based on the mass to charge (m/z) value and the retention time of the ion signals. Ions from both ESI- or ESI+ were merged and imported into the SIMCA-P program (version 14.1) for multivariate analysis. A Principal Components Analysis (PCA) was used as an unsupervised method for data visualization and outlier identification. The pathway enrichment analysis was performed with MetaboAnalyst 5.0 (*33*).

### 4-Hydroxyphenyllactic acid (HPLA) and nitisinone administration in mice

Mice were fed with DL-HPLA (Fisher Scientific, Hampton, NH) (1mg/kg) or the 4-hydroxyphenylpyruvate dioxygenase inhibitor, nitisinone (Sigma-Aldrich, St. Louis, MO) (1mg/kg) with a curved feeding needle. Tick challenge was started 2 hours after being administered HPLA or nitisinone. Tick attachment, tick weight and survival were recorded.

### Statistical analysis

The statistical analysis was performed with Prism 9 software (GraphPad Software, San Diego, CA, USA). To assess the significance of the data on tick weights, a paired t-test was used compare two groups, and an analysis of variance (ANOVA) was used to compare variances across three groups. To compare the tick attachment rate at different time points, a multiple paired t-test was used. The log-rank (Mantel-Cox) test was used to assess survival curves.

## Supporting information

supplemental table 1. metaboloms of sera of guinea pigs (negative ion mode)

supplemental table 2. metaboloms of sera of guinea pigs (postive ion mode)

supplemental table 3. metaboloms of sera of mice (negative ion mode)

supplemental table 4. metaboloms of sera of mice (postive ion mode)

## Author contributions

Conceptualization: YC, EF; Methodology: YC, EF; Investigation: YC, JM, XT, CB, KD, MW; Data analysis: YC; Supervision: EF; Writing— original draft: YC, EF; Writing—review & editing: YC, JM, XT, CB, KD, MW, EF.

## Data and materials availability

All data are available in the main text or the supplementary materials.

